# Multi-metabolomics using imaging mass spectrometry and liquid chromatography-tandem mass spectrometry for spatially characterizing monoterpene indole alkaloids secreted from roots

**DOI:** 10.1101/2021.01.15.426846

**Authors:** Ryo Nakabayashi, Noriko Takeda-Kamiya, Yutaka Yamada, Tetsuya Mori, Mai Uzaki, Takashi Nirasawa, Kiminori Toyooka, Kazuki Saito

## Abstract

Plants release specialized (secondary) metabolites from their roots to communicate with other organisms, including soil microorganisms. The spatial behavior of such metabolites around these roots can help us understand roles for the communication; however, currently they are unclear because soil-based studies are complex. Here, we established a multi-metabolomics approach using imaging mass spectrometry (IMS) and liquid chromatography-tandem mass spectrometry (LC-MS/MS) to spatially assign metabolites under laboratory conditions using agar. In a case study using *Catharanthus roseus*, we showed that 58 nitrogen (N)-containing metabolites are released from the roots into the agar. For the metabolite assignment, we used ^15^N-labeled and nonlabeled LC-MS/MS data, previously reported. Four metabolite ions were identified using authentic standard compounds as derived from monoterpene indole alkaloids (MIAs) such as ajmalicine, catharanthine, serpentine, and yohimbine. An alkaloid network analysis using dot products and spinglass methods characterized five clusters to which the 58 ions belong. The analysis clustered ions from the indolic skeleton-type MIAs to a cluster, suggesting that other communities may represent distinct metabolite groups. For future chemical assignments of the serpentine community, key fragmentation patterns were characterized using the ^15^N-labeled and nonlabeled MS/MS spectra.

---

The development of “omics” technologies has shed light on soil studies^1^. Sequencing technologies have generated huge quantities of microorganisms’ genome/transcriptome data, thereby helping us understand different populations or behaviors in the soil^2^. Metabolome data are key to understanding of how plants communicate with organisms, e.g., how metabolites keep enemies from roots or attract beneficial organisms. Recent studies have shown that specialized metabolites alter rhizosphere microbiota^1,3,4^.

To date, metabolomics approaches have been poorly developed for soil studies. The soil chelates or degrades metabolites, which are released from roots. In addition, metabolite chemical composition and specificity can also exacerbate the development of such approaches. Sequencing technologies have been used to successfully annotate genomes or gene information from sequencing data using huge comprehensive databases; however, it is difficult for metabolomics to adequately assign metabolite information because of poor database resources. Thus, new approaches are required to understand how metabolites function in and around roots and what type of metabolites are released from the roots, even under experimental conditions.

Matrix-assisted laser desorption/ionization-Fourier transform ion cyclotron resonance-imaging mass spectrometry (MALDI-FTICR-IMS) is an important technology for the understanding of the spatial metabolome of plant tissue longitudinal or cross sections with ultra-high resolution and accuracy of ion peaks^5-7^. Matrix reagents are sprayed onto the sections to extract metabolites from surfaces. After this, mixed crystals of the matrix reagent and extracted metabolites are generated on these surfaces. Then, MALDI-FTICR-IMS analysis is conducted on the sections. Spatial signal intensities of metabolites are used for visualization, where higher and lower signal intensities reflect metabolite localization.

Liquid chromatography-tandem mass spectrometry (LC-MS/MS) chemically assigns specialized (secondary) metabolites^8-11^. From analyses, structural information such as retention times, mass to charge (*m/z*) values of precursor and product ions, and fragmentation patterns are generated. Metabolite identification using authentic standard compounds is recommended, but annotation using reported spectra or characterization using deconvoluted spectra is also important in identifying metabolites^12^.

Here, we developed a multi-metabolomics approach using MALDI-FTICR-IMS and LC-MS/MS under experimental conditions, using agar. Even though metabolite behavior in agar is theoretically different to soil, the generation of fundamental data will support field experiments. We used the medicinal plant *Catharanthus roseus* because this plant biosynthesizes monoterpene indole alkaloids (MIAs)^6,13,14^, the functions of which are currently unknown. The workflow comprised: 1) sectioning agar and roots; 2) performing MALDI-FTICR-IMS on sections; 3) performing LC-MS/MS on agar, roots, and leaf samples; and 4) reusing ^15^N-labeled and nonlabeled metabolome data^14^ for the chemical assignment of nitrogen (N)-containing metabolites, including MIAs. From this approach, we characterized 58 N-metabolites including MIAs from roots.

## Experimental section

### Chemicals

Ajmalicine [Sigma-Aldrich Japan (Tokyo, Japan)], catharanthine [Sigma-Aldrich Japan (Tokyo, Japan)], serpentine [FUJIFILM Wako Pure Chemical Corporation, (Osaka, Japan)], and yohimbine [FUJIFILM Wako Pure Chemical Corporation, (Osaka, Japan)] were used in this study.

### Plant materials and growth conditions

*Catharanthus roseus* (Equator White Eye, Sakata Seed Corporation) was used in this study. Fifty ml of Murashige and Skoog agar medium [FUJIFILM Wako Pure Chemical Corporation, (Osaka, Japan)], 8 g/l) was prepared in plant boxes (125 ml, VWR, US). The agar samples were harvested from the boxes in which two plants were grown for three months under a log day (16h day/8h night) condition at 24 °C. As a negative control, a box without the plants was incubated at the same time and was harvested. Leaves and roots of the plants were also harvested from the plants. All the samples were immediately lyophilized at −55 °C. The lyophilized materials were stored at room temperature with silica gel. Four independent samples were analyzed in this study.

### Preparing sections

For the IMS analysis, agar was cut with a razor, embedded with a compound (Surgipath FSC22: Leica Microsystems, Germany) and frozen in a −75 °C acetone bath (Histo-Tek Pino: Sakura Finetek Japan Co.,Ltd., Tokyo, Japan). The frozen sample block was placed on a cryostat specimen disk and was cut with the knife blade until the desired tissue surface appeared. Transfer tape (Adhesive Tape Windows, Leica Microsystems, Germany) was placed on the face of the block to obtain sections (each with a thickness of 20 μm) in the CM3050S cryostat (Leica Microsystems, Germany). Two sections were transferred to conductive Cu tape (double-sided, No. 796) (TERAOKA SEISAKUSHO, Co. Ltd.) on a glass slide (ITO coating, Bruker Daltonik GmbH). The section on the glass slide was freeze-dried overnight at −30°C in the cryostat.

### MALDI-FTICR-IMS analysis

A 2,5-dihydroxybenzoic acid (DHB) matrix solution (15 mg/mL in 90% ACN, 0.1% TFA) was sprayed on the prepared section that was on the glass slide using TM-sprayer (HTX TECHNOLOGIES, LLC) running modified parameters (nozzle temperature, 60 °C; pump device, LC pump; plate rate 0.125 ml/min; z-arm velocity, 700 mm/min; number of passes, 14; moving pattern, CC; track spacing, 3 mm) for a wet condition. The freeze-dried section with the matrix was analyzed in the FTICR−MS SolariX 7.0 T instrument. The MALDI parameters are as follows: geometry, MTP 384 ground steel; plate offset, 100.0 V; deflector plate, 200.0 V; laser power, 40.0%; laser shots, 100; frequency, 2000 Hz; laser focus, small; raster width, 30 μm. The MS conditions were as follows: mass range, *m/z* 100.33-500.00; average scan, 1; accumulation, 1.000 s; polarity, positive; source quench, on; API high voltage, on; resolving power, 66,000 at 400 *m/z*; transient length, 0.4893 s; Mode (data storage: save reduced profile spectrum, on; reduced profile spectrum peak list, on; data reduction, 97%; auto calibration: online calibration, on; mode, single; reference mass, *m/z* 362.926338); API Source (API source: source, ESI; capillary, 4500 V, end plate offset, −500; source gas tune: nebulizer, 1.0 bar; dry gas, 4.0 l/min; dry temperature, 180 °C); Ion Transfer (source optics: capillary exit, 180 V; detector plate, 200 V; funnel 1, 150 V; skimmer 1, 10 V; funnel RF amplitude, 120 Vpp; octopole: frequency, 5 MHz; RF amplitude, 250 Vpp; quadrupole: Q1 mass, 100.0 *m/z*; collision cell: collision voltage, −3.0 V; DC extract bias, 0.3 V; RF frequency, 2 MHz; collision RF amplitude, 600.0 Vpp; transfer optics: time of flight, 0.400 ms; frequency, 6 MHz; RF amplitude, 200.0 Vpp); Analyzer (infinity cell: transfer exit lens, −20.0 V; analyzer entrance, −10.0 V; side kick, 0.0 V; side kick offset, −1.0 V; front trap plate, 0.700 V; back trap plate, 0.550 V; sweep excitation power, 12.0%; multiple cell accumulations: ICR cell fills, 1).

### Visualizing IMS data

Visualization was performed using SCiLS Lab software version 2019c (Bruker Daltonik GmbH, Bremen, Germany).

### Metabolite extraction for LC-MS/MS analysis

The freeze-dried agar samples (each, ∼700 mg DW) were extracted with 4.8 ml of 80% MeOH using a mixer (EYELA CUTE MIXER CM-1000) at 1,500 rpm for 7 min. After centrifugation with the following conditions (force, 13,000 *g*; running time, 10 min; and temperature, 4 °C), 3.6 ml of supernatant was dried up. Extracts were resolved with 3 ml of 2.5% MeOH. Extraction solvents were applied to HLB cartridges (3 cc, Waters) after equilibration according to the protocol. The cartridges were washed with 3 ml of 0.1% acetic acid twice, and then were extracted with 3 ml of 90% MeOH. The elution solvents were concentrated and dried up completely. The extracts were resolved with 100 µL of 80% MeOH including 2.5 µM lidocaine and then were filtered through Ultrafree MC centrifugal filter, Millipore).

Extraction of leaf and root samples was performed according to the previous research^14^.

### LC-MS/MS analysis

The analysis was performed according to the previous research^14^.

### Data analysis for N-metabolites

The ^15^N-labeled and nonlabeled metabolome data acquired in the previous study^14^ were used for assignment of N-metabolites. Using the data, ions derived from N-metabolites were assigned to the current data with the following conditions (retention time, ± 0.2 min; *m/z* value, ± 0.01) (**Table S1**).

### Alkaloid network analysis

MS/MS similarities were calculated using dot product algorithm^12^. The similarities more than 0.8 was used for the network. The ions were clustered using spinglass algorithm of R (https://igraph.org/r/doc/cluster_spinglass.html). Parameters on gamma and spins were changed to 1.5 and 200, respectively. Parameters on the network are available in **SI Data S1**. Visualization of the network was performed using the PlaSMA database (http://plasma.riken.jp/).

## Results and discussion

Plants were grown in plant boxes for three months. Agar around the roots was cut away from the box (**Figure 1a**). Cross sections, including the agar and the roots, were prepared using a cryostat. Sections were then placed on conductive tape using transfer tape^15^. As shown in **Figure 1b**, the root was placed in the center. To consider spreading out metabolites from the root during spraying a matrix reagent, the TM sprayer was selected for a much shorter spraying time. The sprayer generated fine-grained mixed crystals of the reagent and metabolites from the root. A larger crystal generated incorrect localization in highly spatial resolution analysis. The spatial resolution was set at 30 µm. IMS analysis was performed in approximately 3 mm^2^ areas to detect metabolites. Thus, ions from root metabolites were detected at approximately 500 µm around the root (**Figure 1c**).

**Figure 1.**
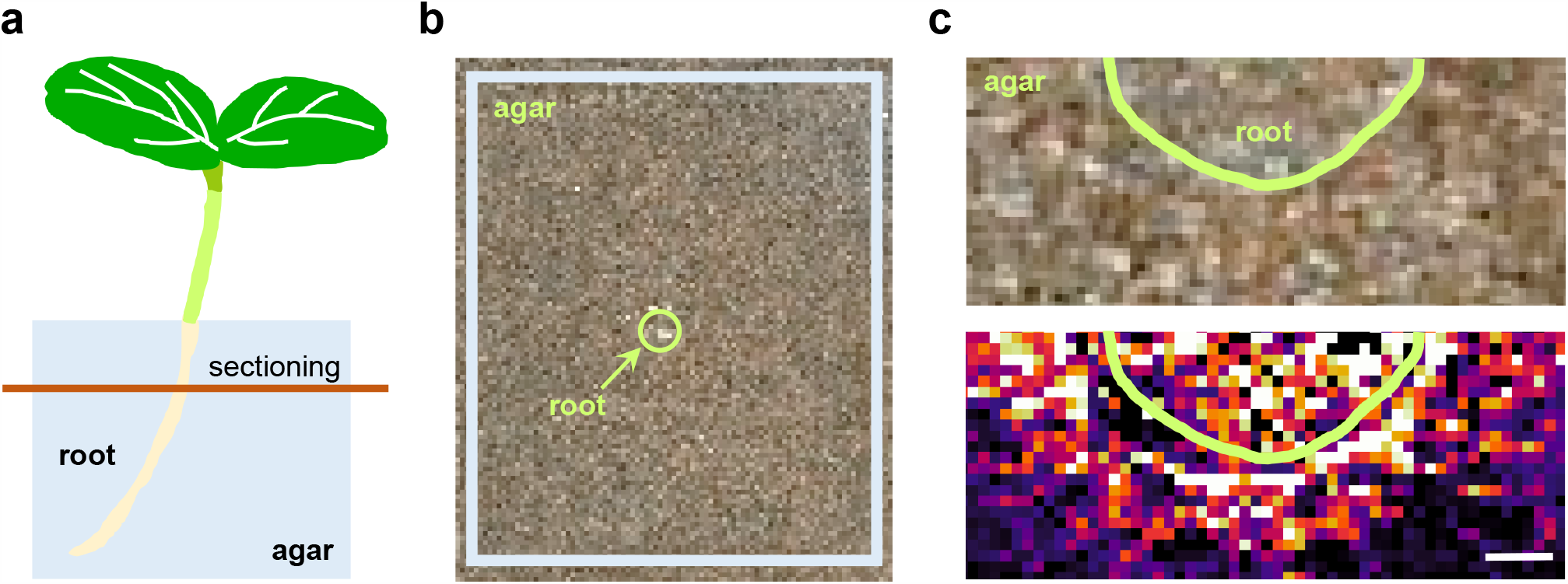
Spatial metabolomics using MALDI-FTICR-IMS. (a) Illustration about the samples for preparing sections. The root was cut out with the agar. (b) The agar including the root was sectioned together and was transferred onto a double-sided conductive tape placed on a glass. (c) Upper. Area analyzed. Lower. Visualization of serpentine ion at *m/z* 349.15464 ± 0.001. Green line indicates the side of the root. Bar means 200 µm.

To chemically assign MIAs, agar samples were analyzed using LC-MS/MS together with leaf and root samples. In addition, agar, where plants were never grown, was analyzed as a negative control. A comparative analysis using both agar samples suggested that large quantities of MIA metabolites are secreted from this. Ions from the LC-MS/MS data were matched using conditions (retention time, ± 0.2 min; *m/z* value, ± 0.01) to the previous data of ^15^N-labeled and nonlabeled plants^14^. After this, ions detected by IMS and LC-MS/MS were paired with a filter condition (*m/z* value, ± 0.005). Finally, LC-MS/MS-based 58 ions were characterized as N-metabolites (**Supporting Information (SI) Table S1**). In consideration of overlapping *m/z* values, of which, 28 ions could be visualized using the IMS data (**SI Figures S1**).

A heat map indicated relative signal intensity patterns of these ions (**Figure 2a**). Using authentic standard compounds, four ions were identified as derived from ajmalicine, catharanthine, serpentine, and yohimbine. Including those, 16 ions were derived from MIAs as having an indolic skeleton, which were characterized in the previous study. Interestingly, the relative signal intensity of the 16 ions was higher. This suggested that this plant releases indolic skeleton-type MIAs from the roots in large quantities. To characterize structural features, an alkaloid network analysis was performed using both MS/MS similarity network and spinglass cluster detection algorithms (**SI Data S1**). The analysis generated five clusters as upper layer and MS/MS similarity network as lower (**Figure 2b, SI Table S1**). The clusters were generated based on MS/MS similarities. When nodes in a cluster had a similarity to others in another cluster, both clusters were connected as shown (**Figure 2b, upper**). Interestingly, ions derived from identified MIAs were included in cluster 1, suggesting the cluster represented indolic skeleton-type MIAs. The ion derived from serpentine was included in cluster 3. All ions except for the serpentine ion were uncharacterized (**Figure 2b, lower**). Because their MS/MS patterns were similar to each other, a deconvoluted serpentine pattern will be useful for future analysis. Here, a fragmentation pattern was elucidated using ^15^N-labeled and nonlabeled MS/MS spectra (**Figure 2c**). By considering mass shifts from labeling, we successfully characterized fragmented bonds leading to substructures, which will elucidate whole structures of other cluster members.

**Figure 2.**
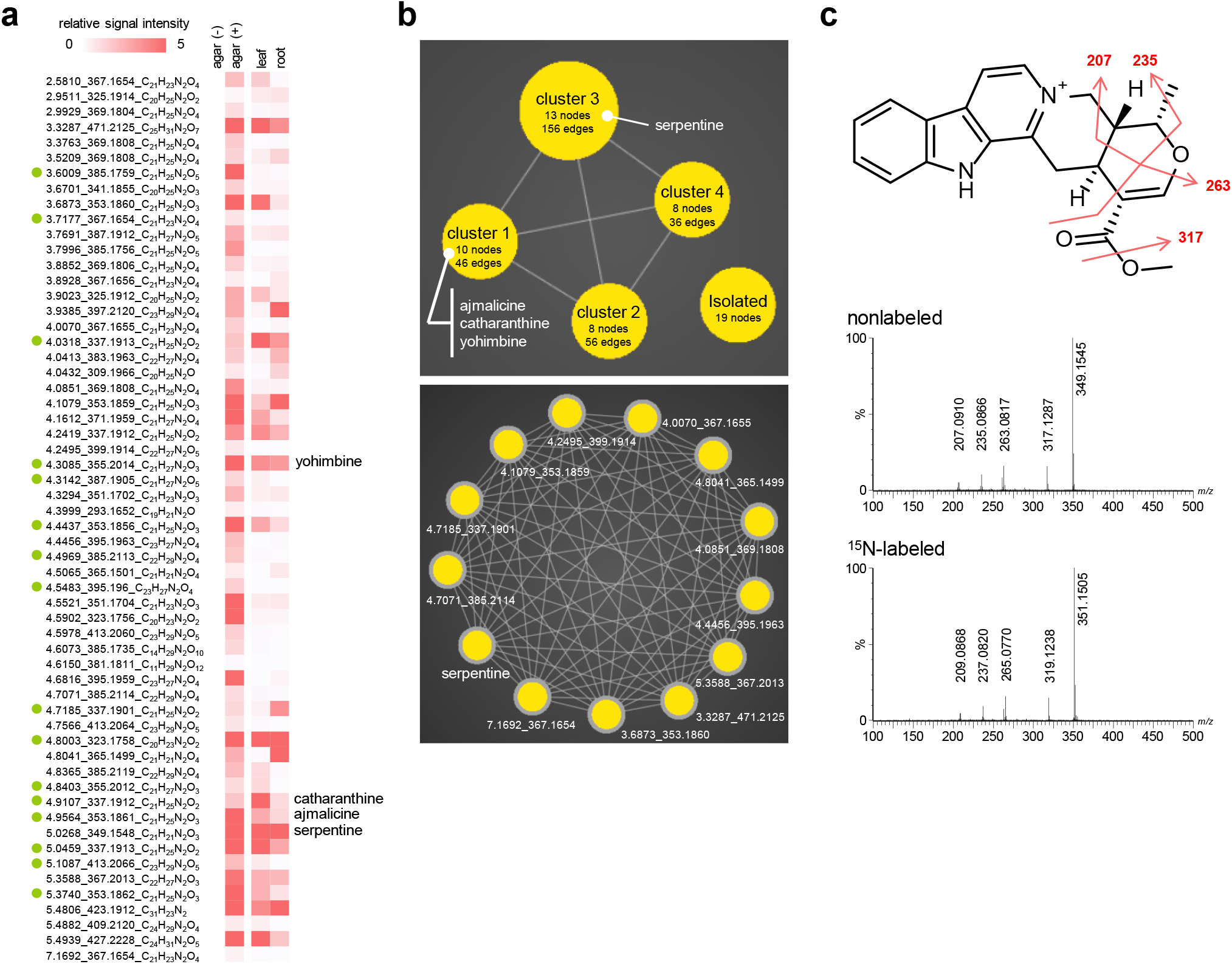
Chemical assignment of N-containing metabolites including MIAs. (a) Heat map using the relative signal intensity of the 58 ions. Light green circle indicates indolic type of MIAs assigned in the previous study. Their retention time and *m/z* value was evaluated with the previous results. (b) Upper. An alkaloid network analysis using MS/MS similarities. In this study, five clusters were obtained. The isolated cluster summarized that ions have less than 0.8 or none similarity to others. Lower. the members of cluster 3. Tag indicates retention time and *m/z* value. (c) Fragmentation analysis of serpentine. Upper. Nonlabeled MS/MS spectrum. Lower. ^15^N-labeled MS/MS spectrum. The numbers indicate *m/z* value of the nonlabeled product ions. MS/MS spectrum of the authentic standard compound was matched to the nonlabeled one. The patterns were considered on the basis of the mass shift due to the ^15^N labeling.

Plants secrete various metabolites from their roots. Primary metabolites such as carbohydrates and organic acids and specialized metabolites are also secreted, including glucosinolates from *Arabidopsis thaliana*, isoflavones from *Glycine max*, and momilactones from *Oryza sativa*^16, 17^. Secreted metabolites play various roles, e.g., nutrient uptake^18^, defense against pathogens and/or other plants^19^, and the regulation of rhizosphere microorganisms^19^. Some studies reported on root exudates containing MIAs; the hairy roots of Catharanthus roseus secrete several MIAs including ajmalicine, serpentine, and catharanthine into the medium^20,21^. Interestingly, endophytic infection induces ajmalicine and serpentine biosynthesis in roots of *C. roseus*, suggesting that these MIAs have roles in rhizosphere interactions with microorganisms^22^.

## Conclusions

In this study, we established the multi-metabolomics approach using MALDI-FTICR-IMS and LC-MS/MS to profile MIAs secreted from *Catharanthus roseus* roots under experimental conditions using agar. This approach identified 58 N-containing metabolites from the roots. Although the functions of these metabolites are still unknown, these data will facilitate in more comprehensive soil/root studies. Four MIAs—ajmalicine, catharanthine, serpentine, and yohimbine—were identified using authentic standard compounds. Applying the MIAs to the soil characterizes the alternation of the microbiota. A comparative analysis using mutants lacking MIAs (using genome editing technologies) will uncover functional profiles and relationships with microbiota. Combining this approach with other omics approaches will help us understand how this plants alter soil microbiota via MIAs.

## Associated Contents

### Supporting Information

The Supporting Information is available free of charge at online.

Figure S1. Visualization of the ions detected in the MALDI-FTICR-IMS and LC-MS/MS analyses.

Table S1. Table S1. Detected ions in the multi-metabolomics. Data S1. Information of the alkaloid network analysis

## Author Contributions

R.N. designed the research. R.N., N.T.-K., T.M., T. N., and K.T., prepared the samples. R.N. and T.M. acquired and analyzed the metabolome data. Y. Y. performed the alkaloid network analysis. R.N., M. U., and K.S. discussed the research. R. N. wrote the manuscript.

## Notes

The authors declare no competing financial interest.

