## Supplementary material for "Multi-metabolomics using imaging mass spectrometry and liquid chromatography-tandem mass spectrometry for spatially characterizing monoterpene indole alkaloids secreted from roots": Figure S1

Content:
**SI Figure S1.** Visualization of the ions detected in the MALDI-FTICR-IMS and LC-MS/MS analyses.

### Figure S1

*m/z* 293.164

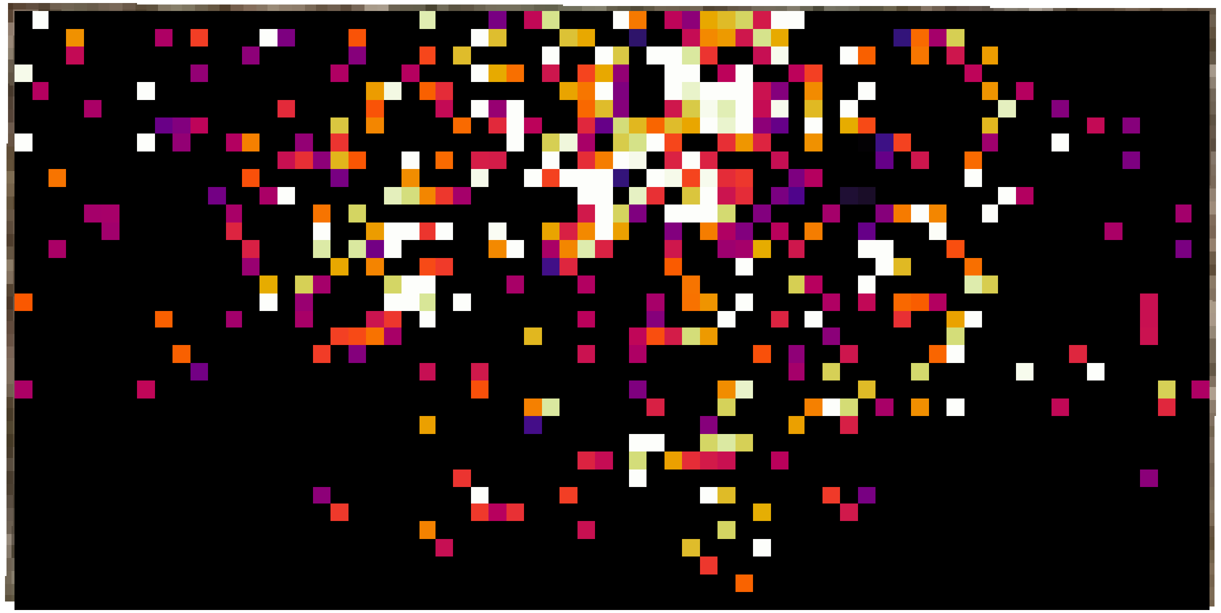

#### Figure S1 (continued)

*m/z* 309.196

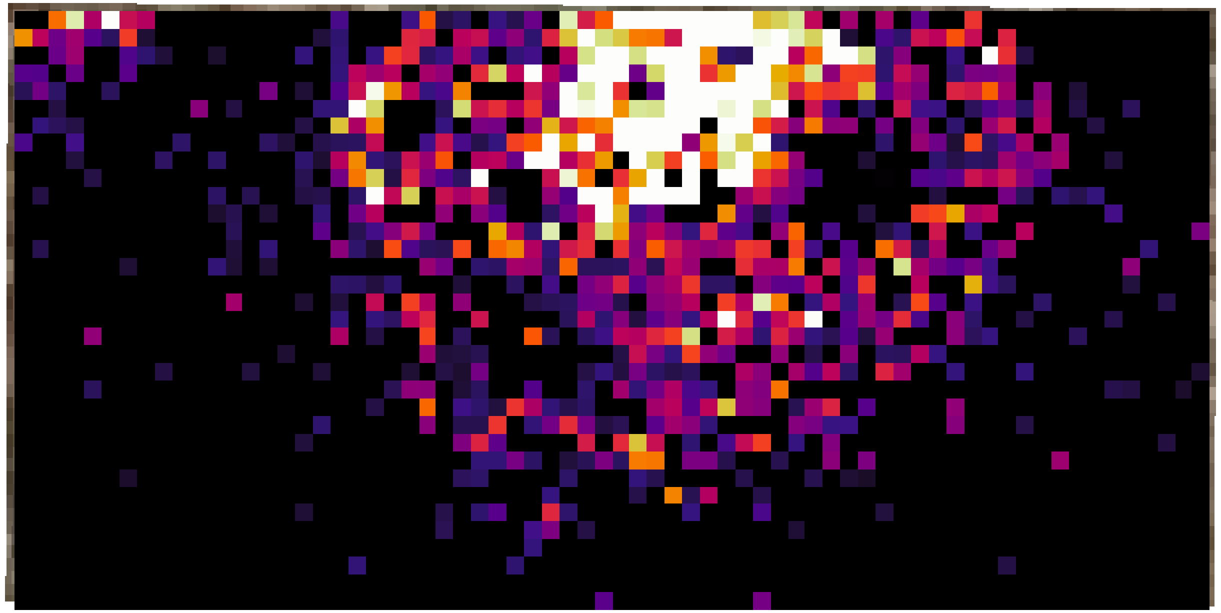

### Figure S1 (continued)

*m/z* 323.175

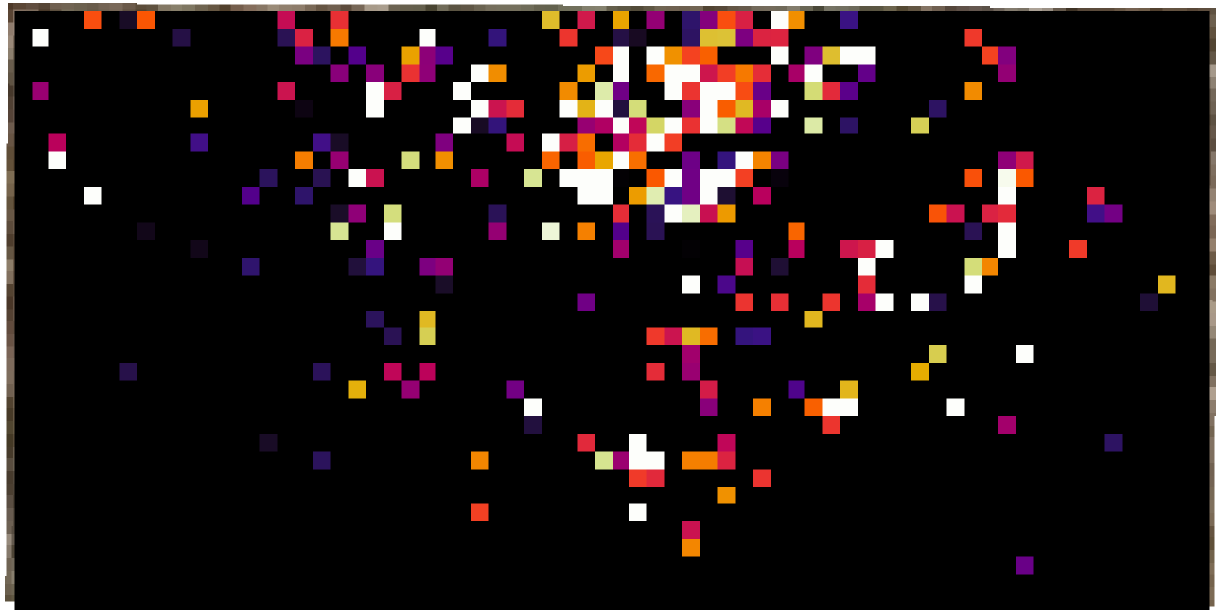

### Figure S1 (continued)

*m/z* 367.165

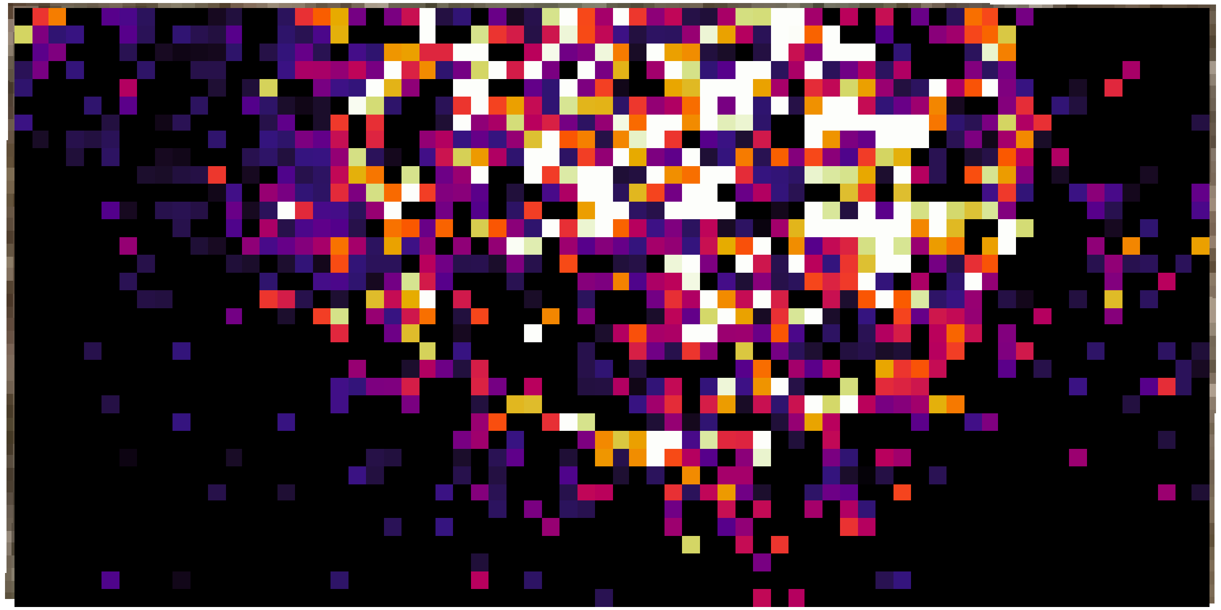

#### Figure S1 (continued)

*m/z* 337.191

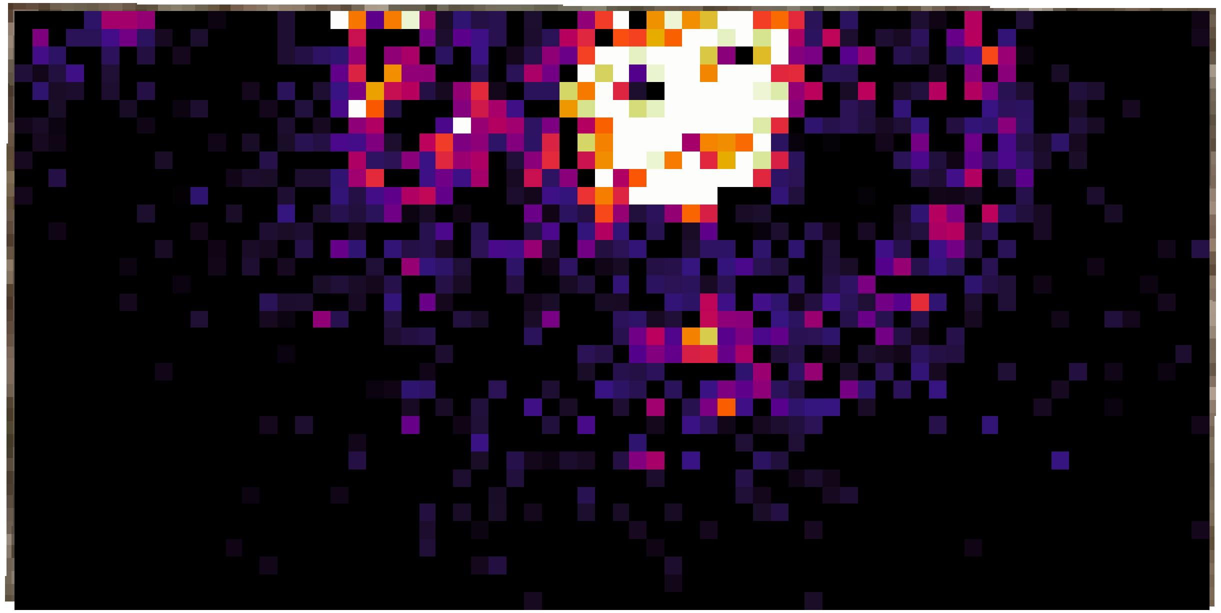

**Figure S1 (continued)**

*m/z* 341.185

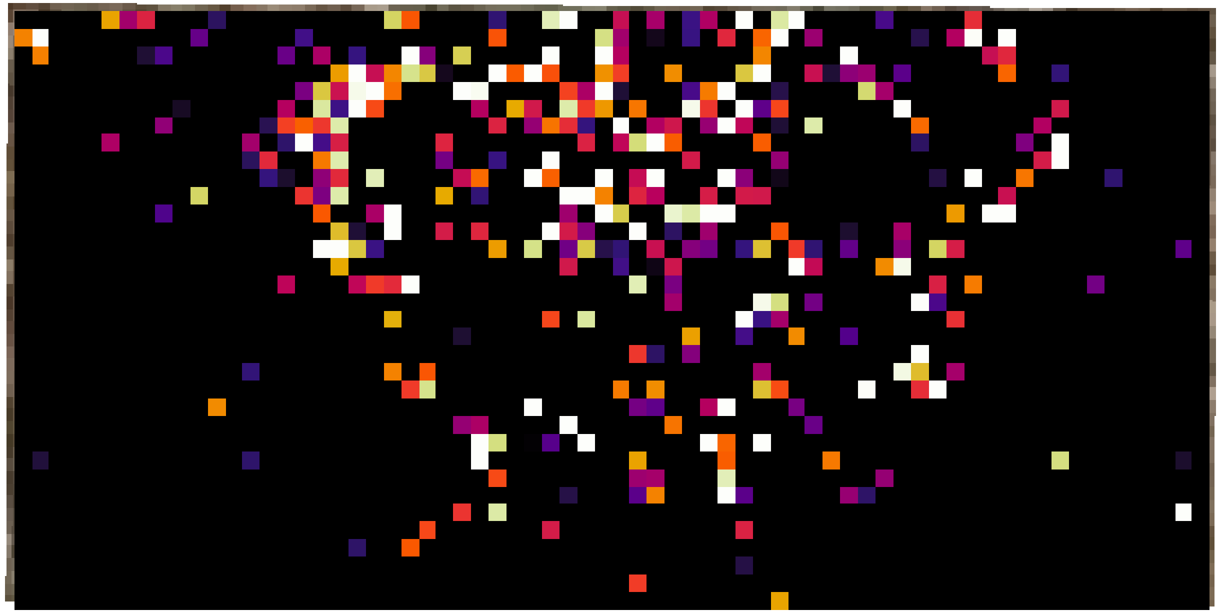

**Figure S1 (continued)**

*m/z* 349.154

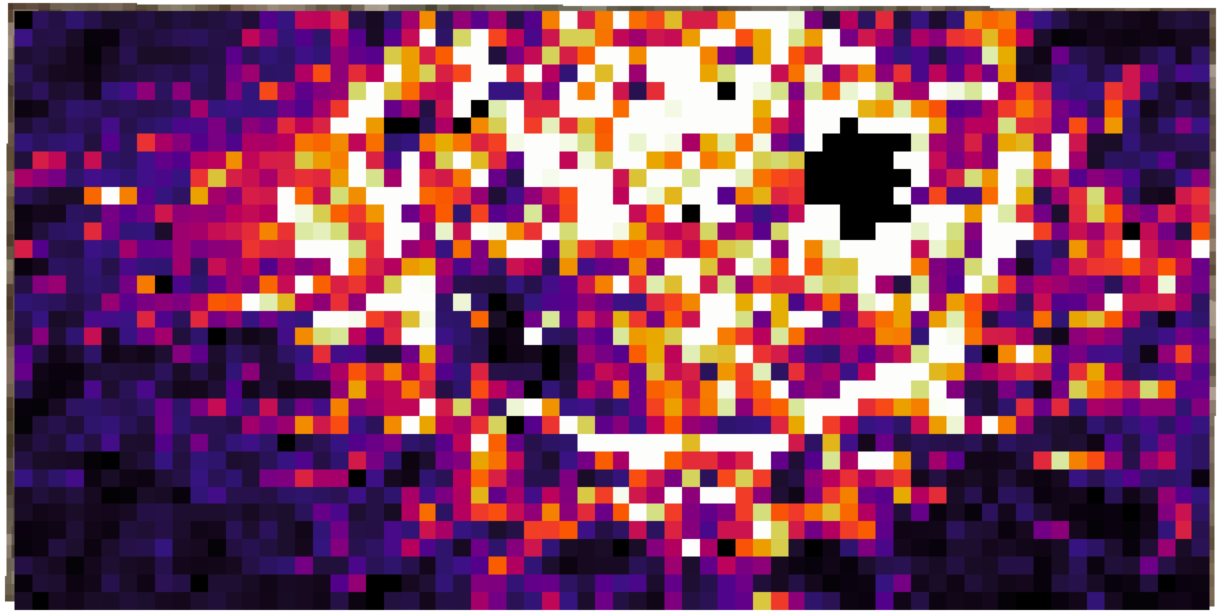

#### Figure S1 (continued)

*m/z* 351.170

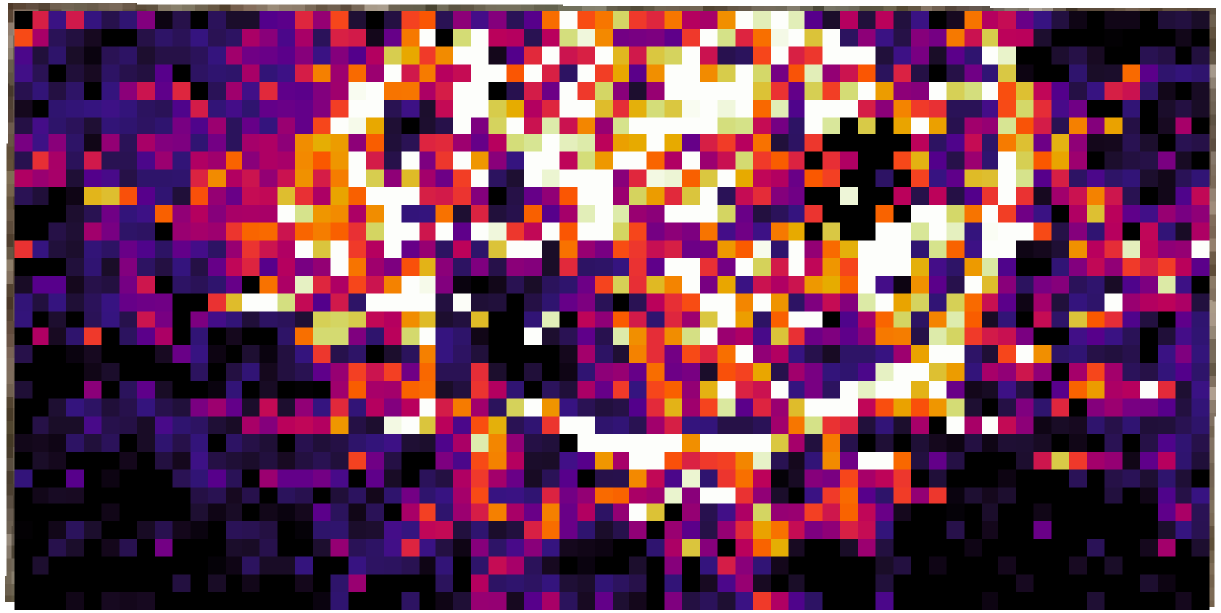

### Figure S1 (continued)

*m/z* 353.185

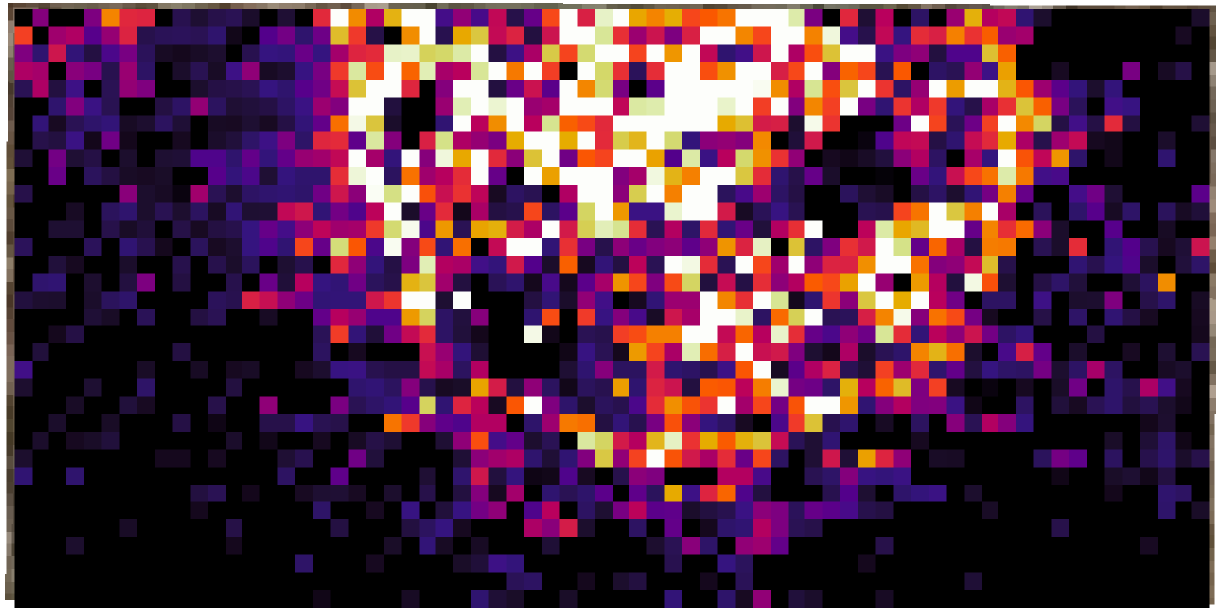

#### Figure S1 (continued)

*m/z* 355.201

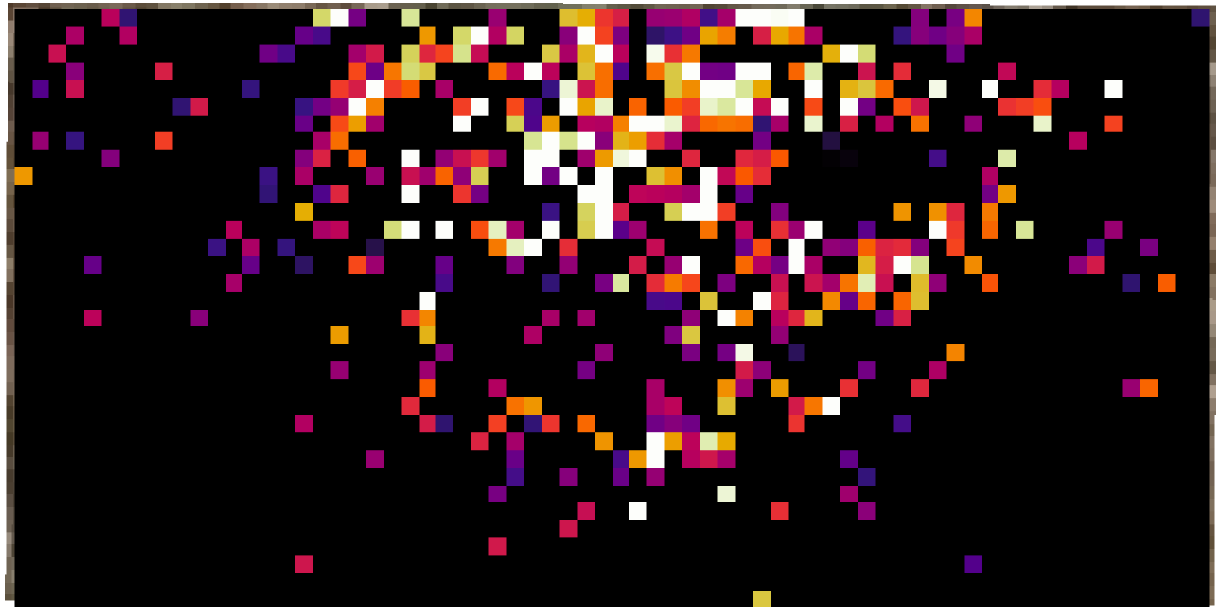

**Figure S1 (continued)**

*m/z* 365.149

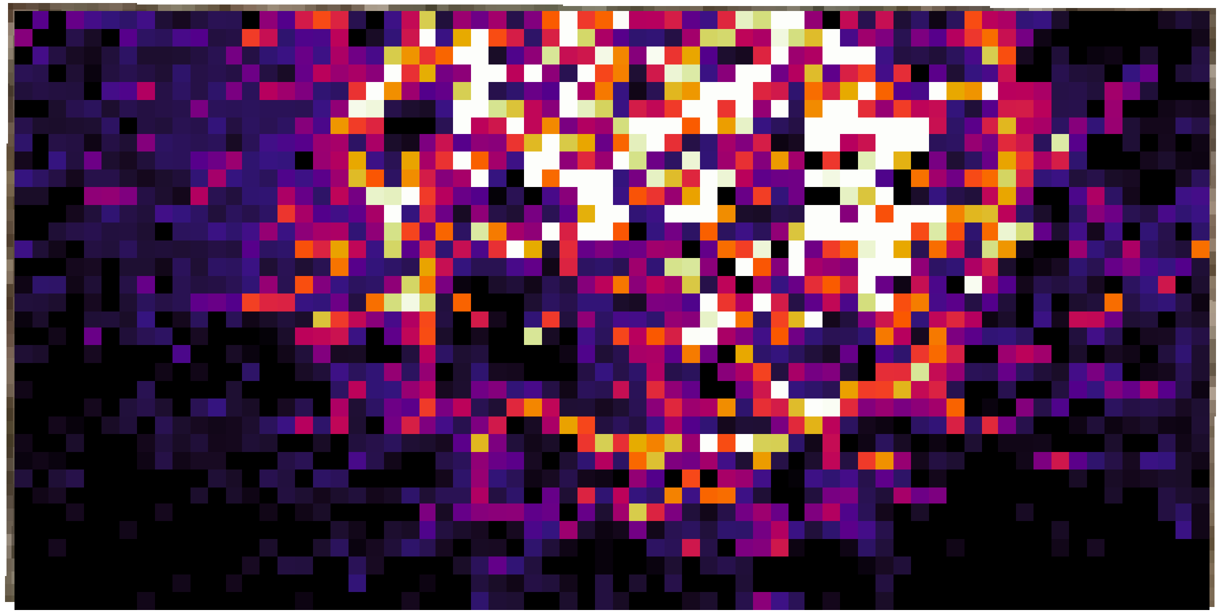

**Figure S1 (continued)**

*m/z* 367.165

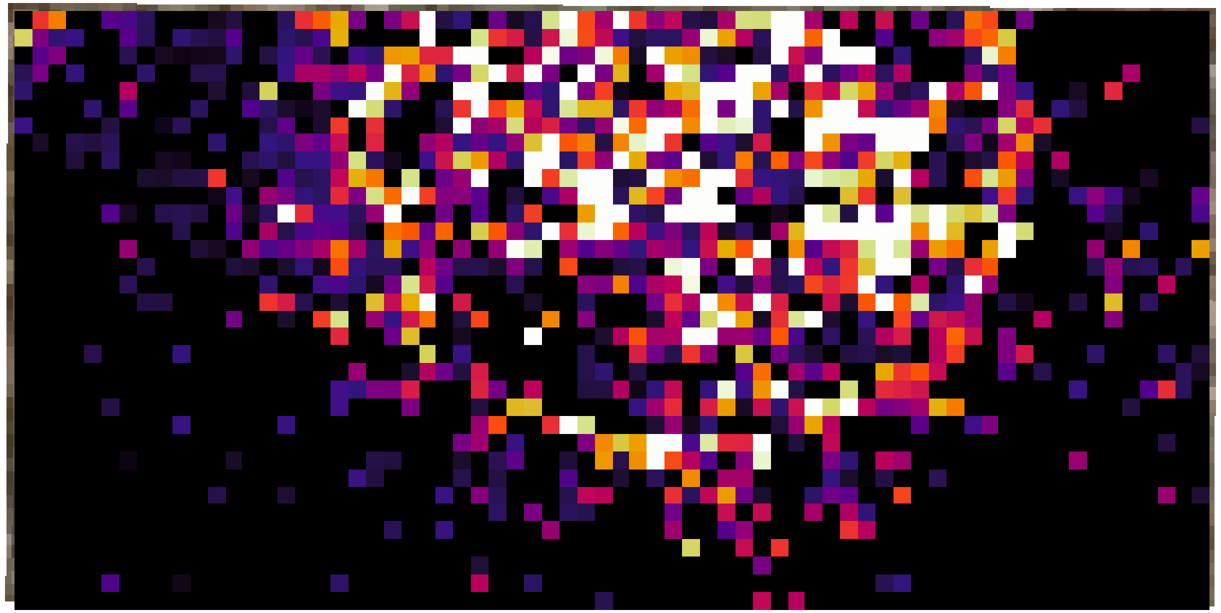

### Figure S1 (continued)

*m/z* 367.201

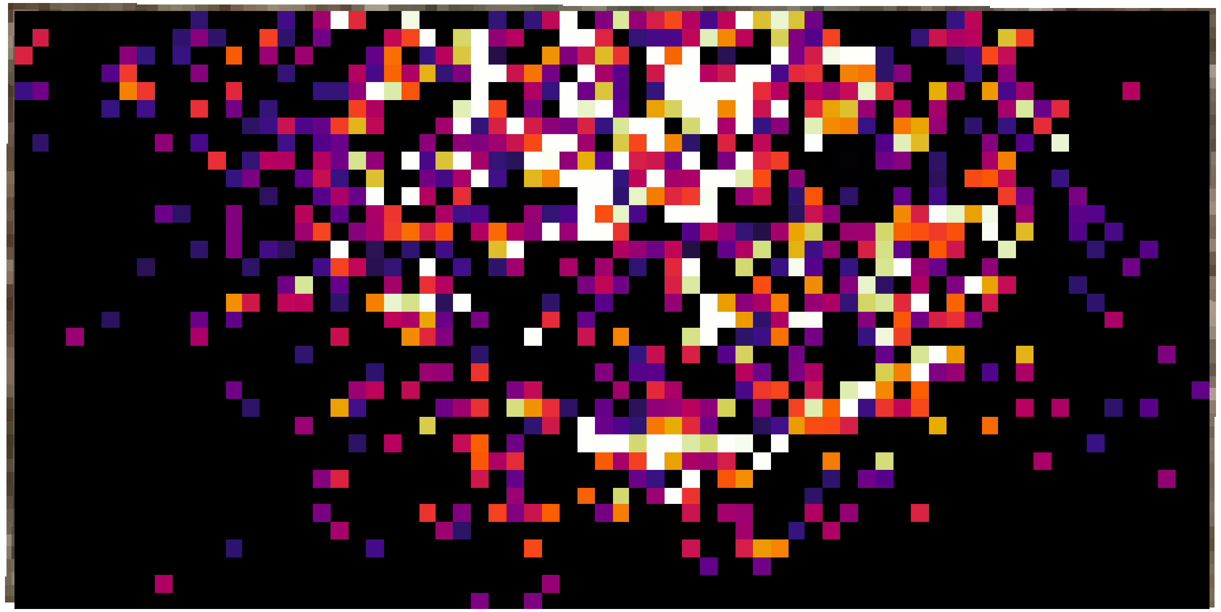

#### Figure S1 (continued)

*m/z* 369.181

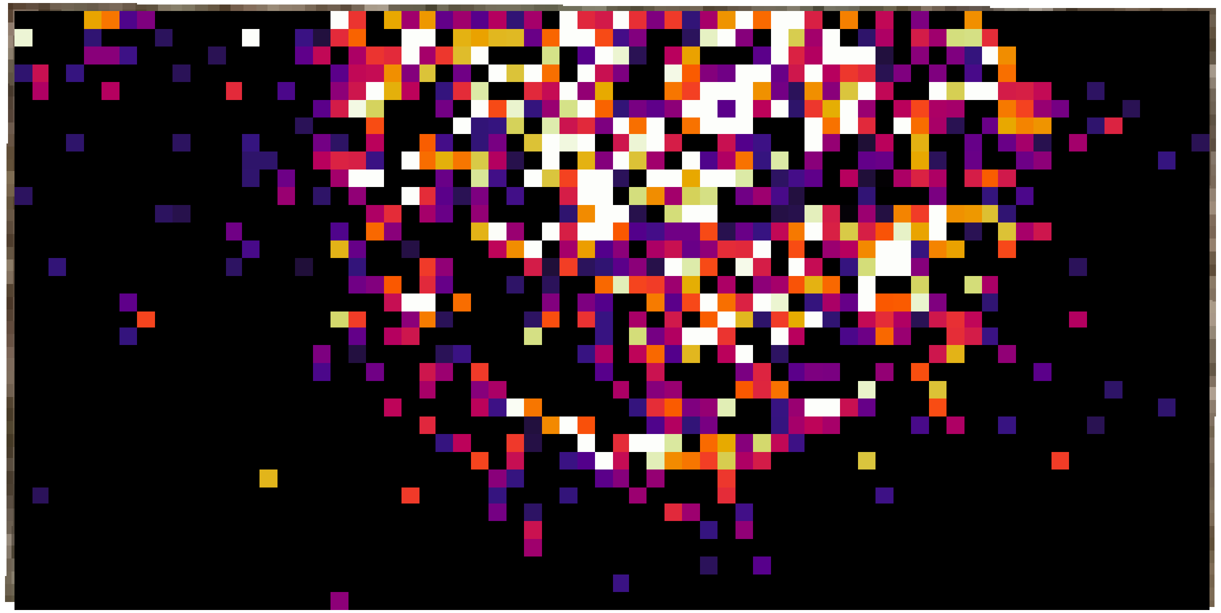

**Figure S1 (continued)**

*m/z* 371.196

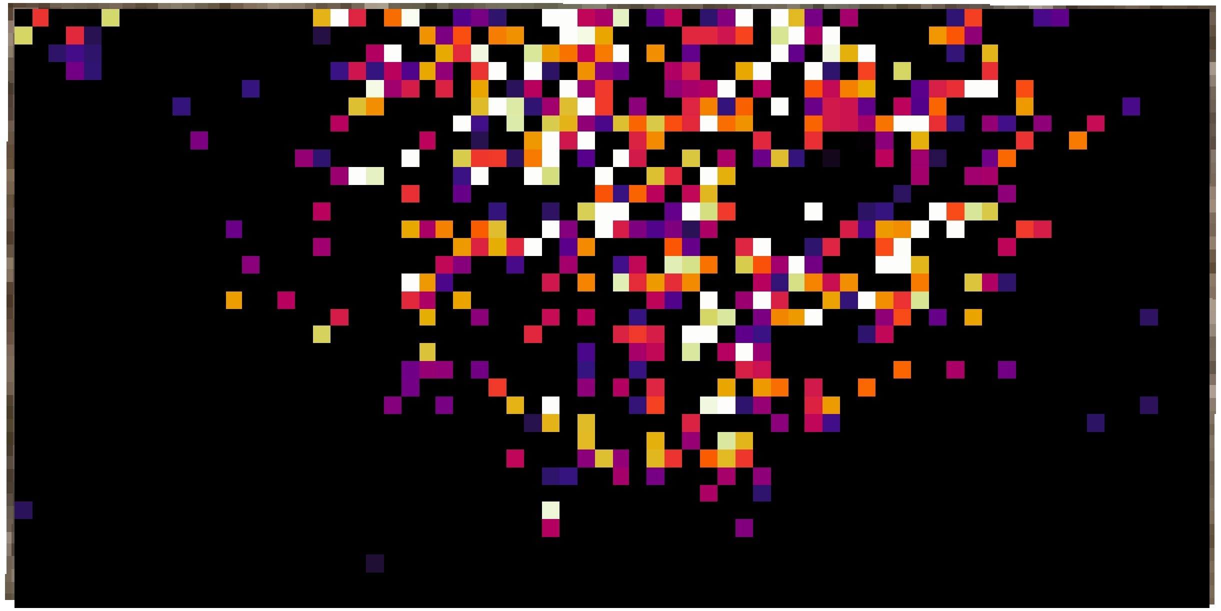

**Figure S1 (continued)**

*m/z* 381.108

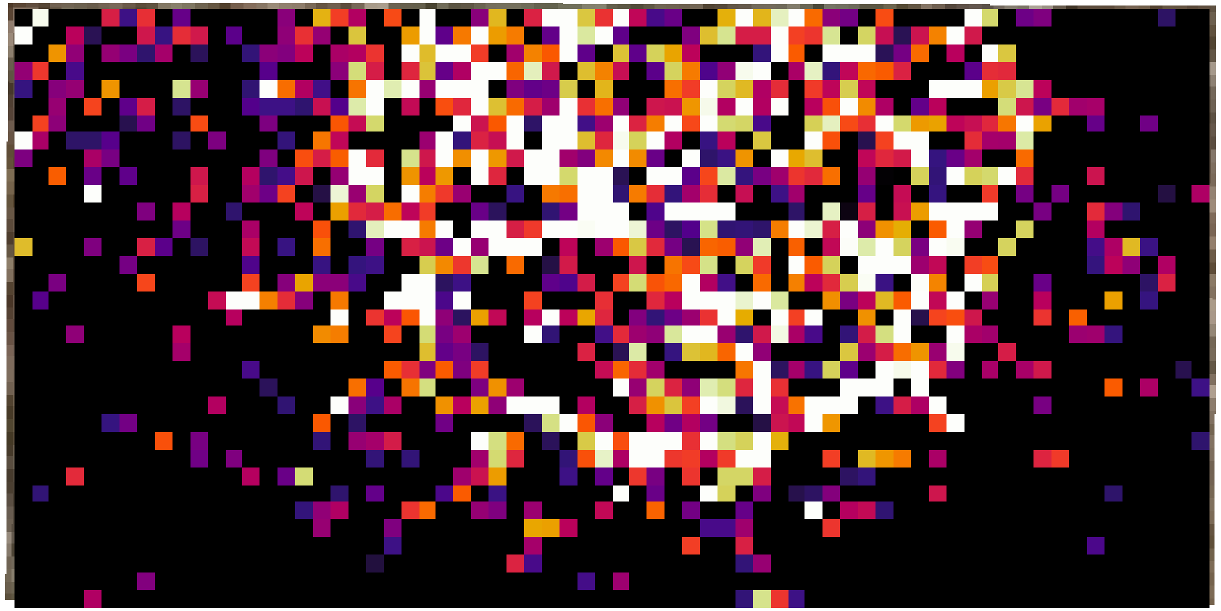

**Figure S1 (continued)**

*m/z* 383.196

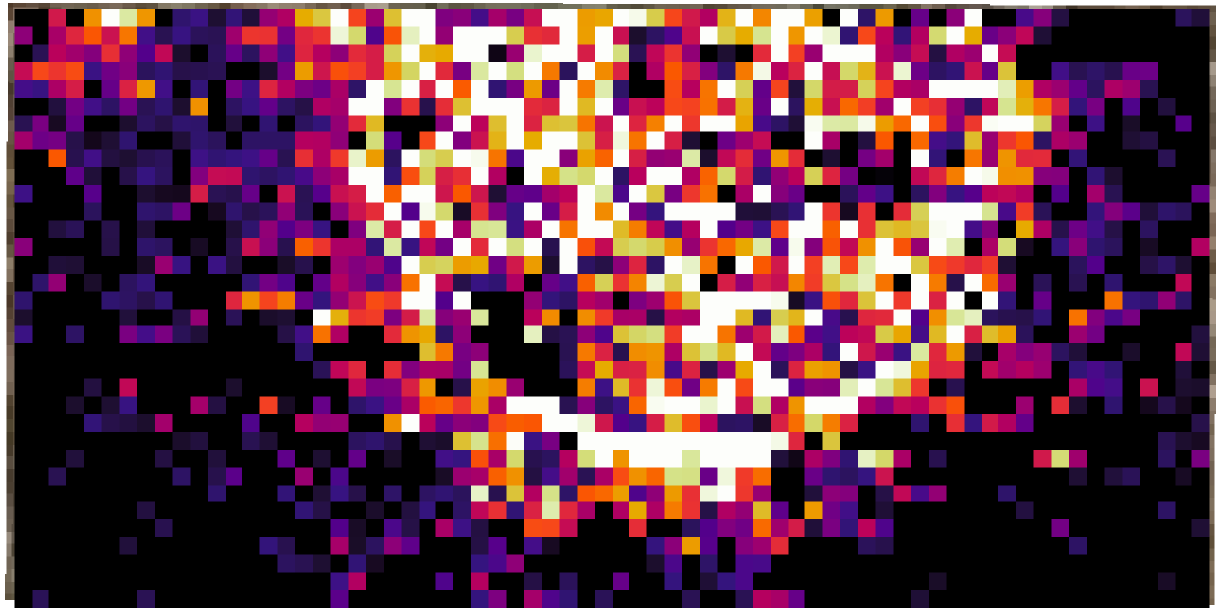

### Figure S1 (continued)

*m/z* 385.175

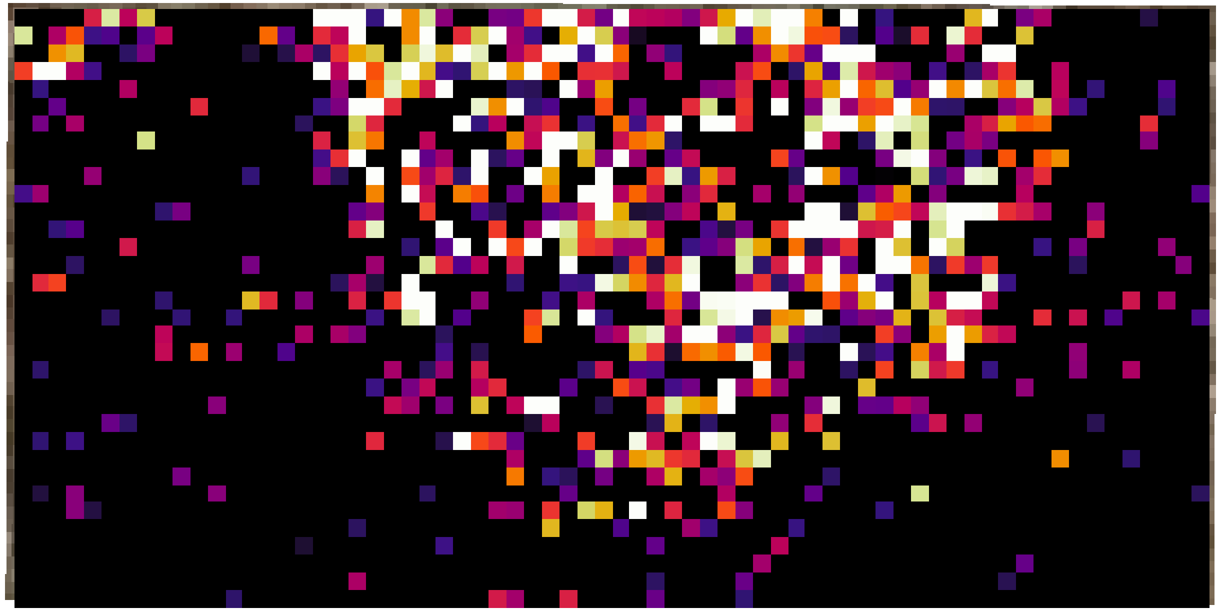

### Figure S1 (continued)

*m/z* 385.211

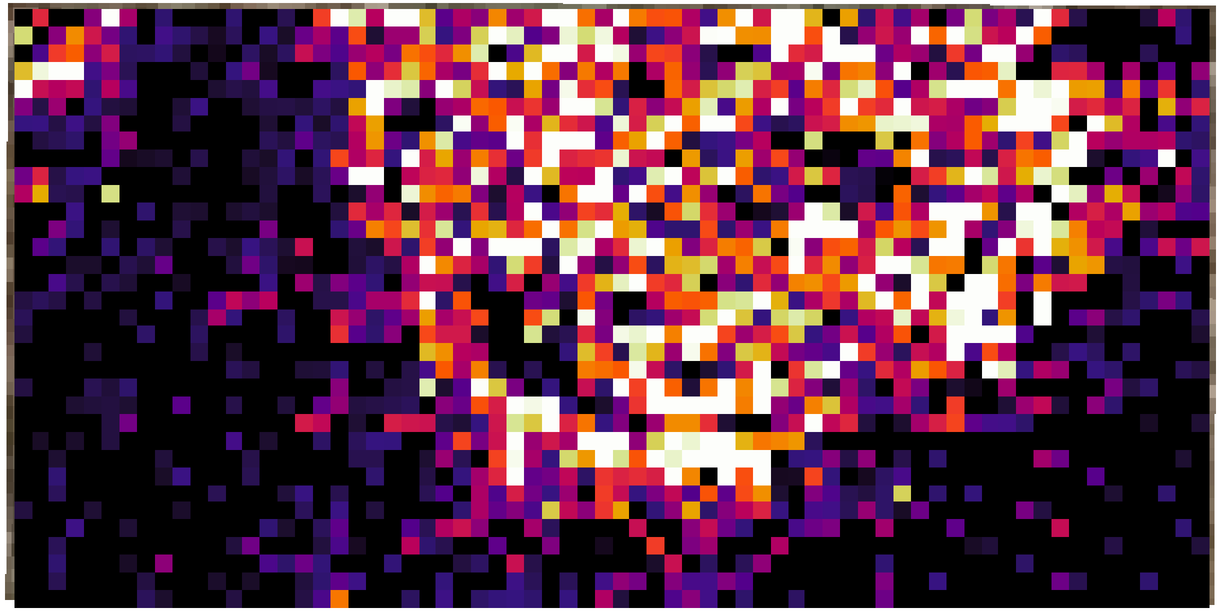

### Figure S1 (continued)

*m/z* 387.191

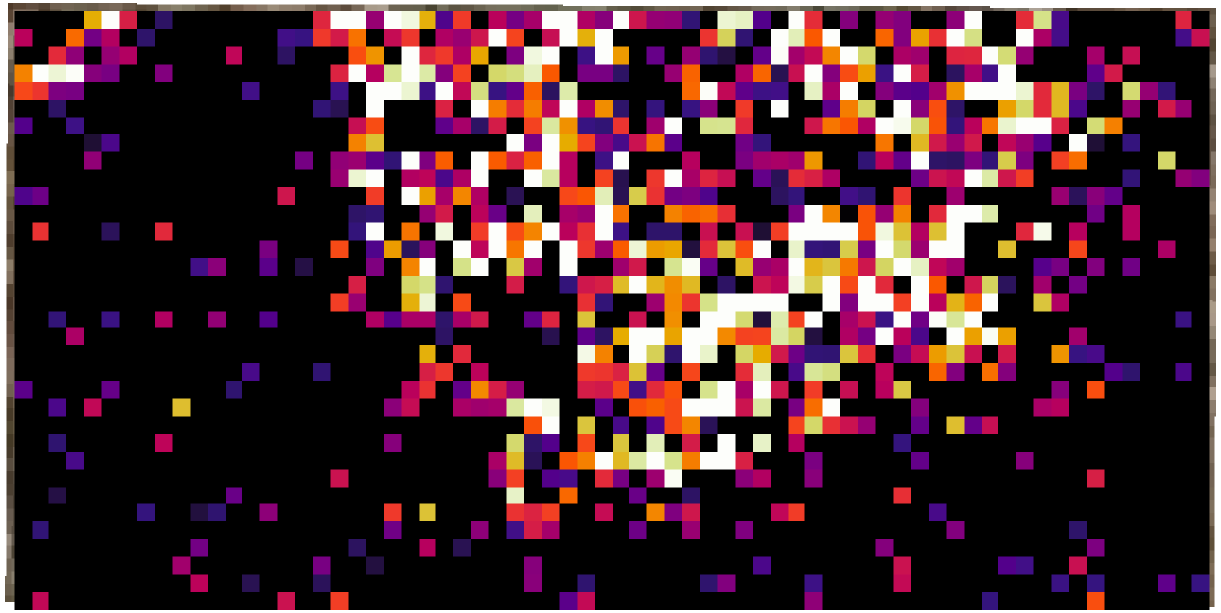

**Figure S1 (continued)**

*m/z* 395.196

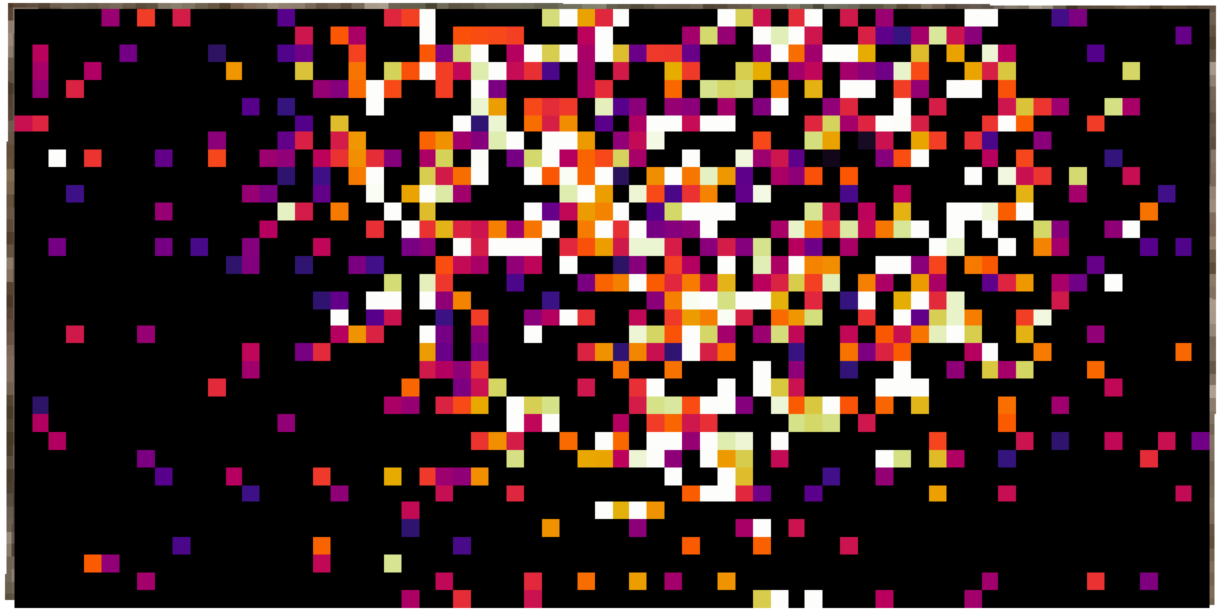

### Figure S1 (continued)

*m/z* 397.212

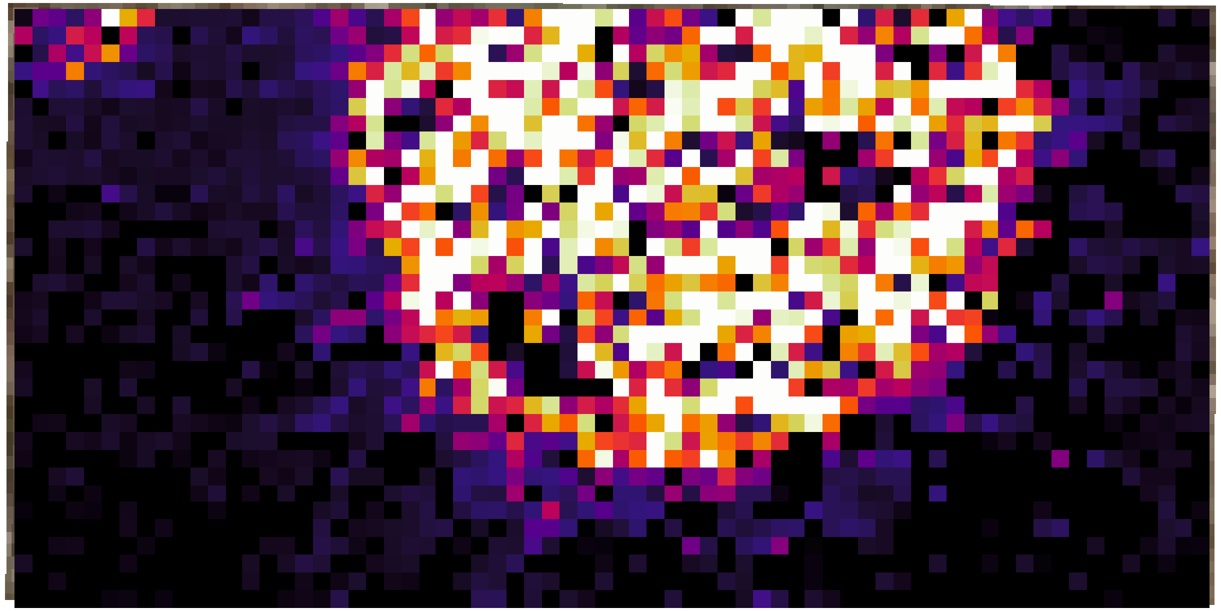

**Figure S1 (continued)**

*m/z* 399.191

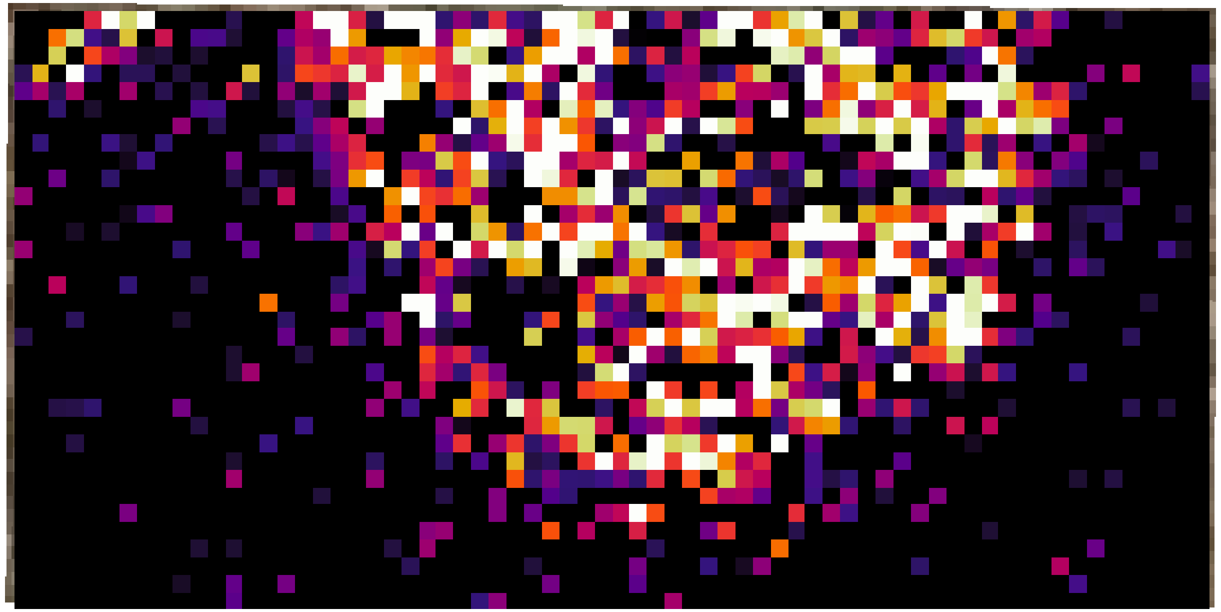

**Figure S1 (continued)**

*m/z* 409.211

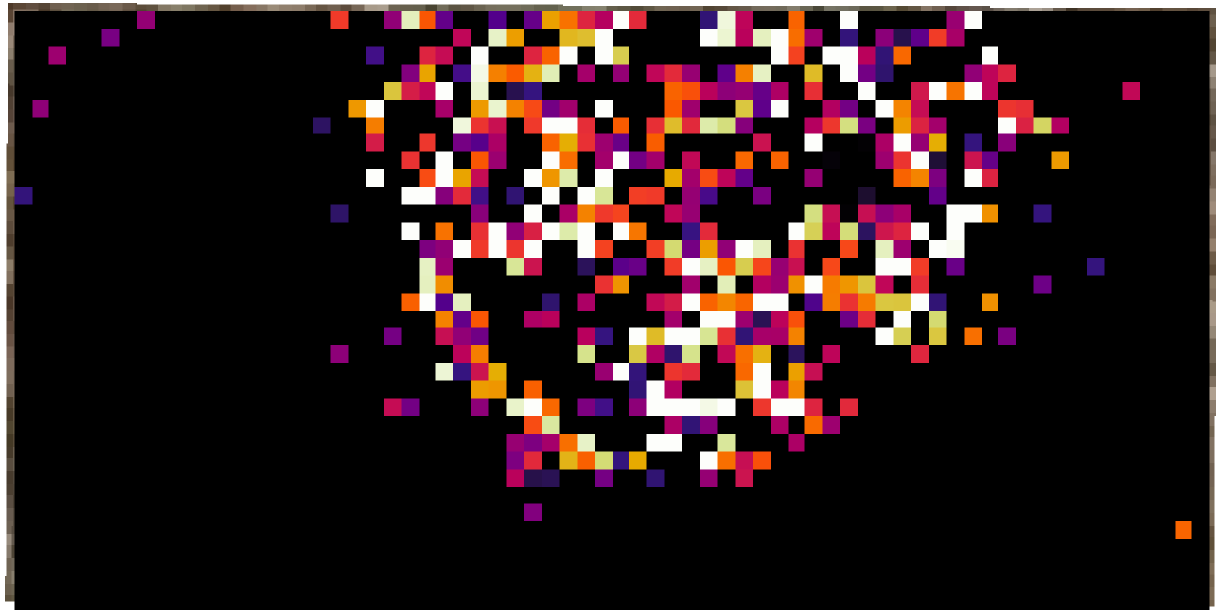

**Figure S1 (continued)**

*m/z* 413.207

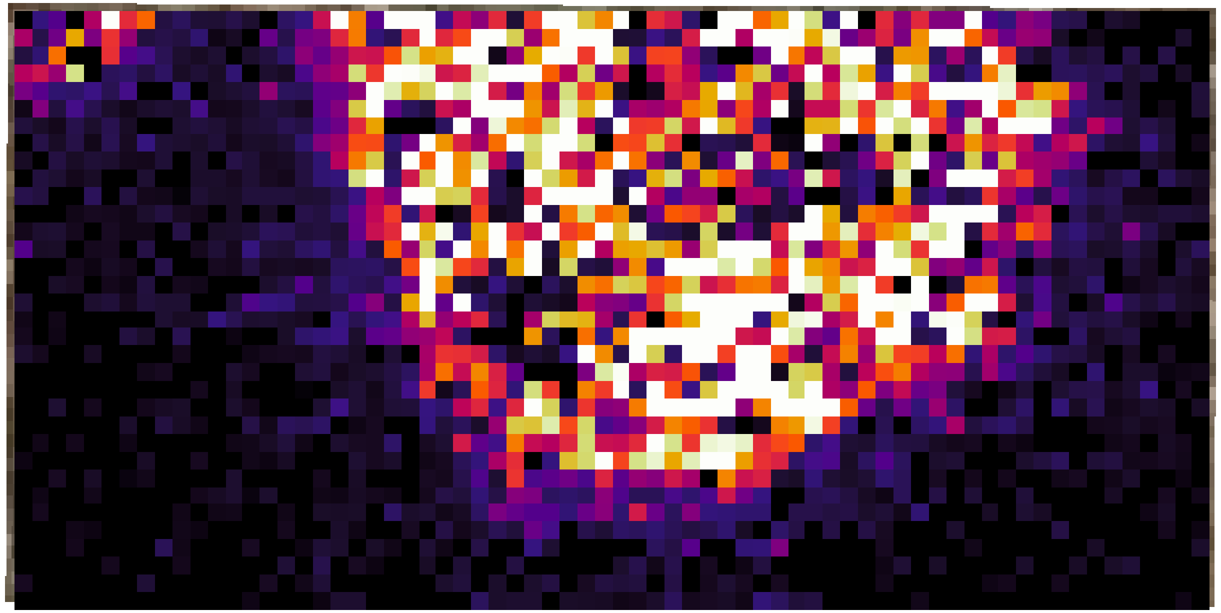

**Figure S1 (continued)**

*m/z* 423.191

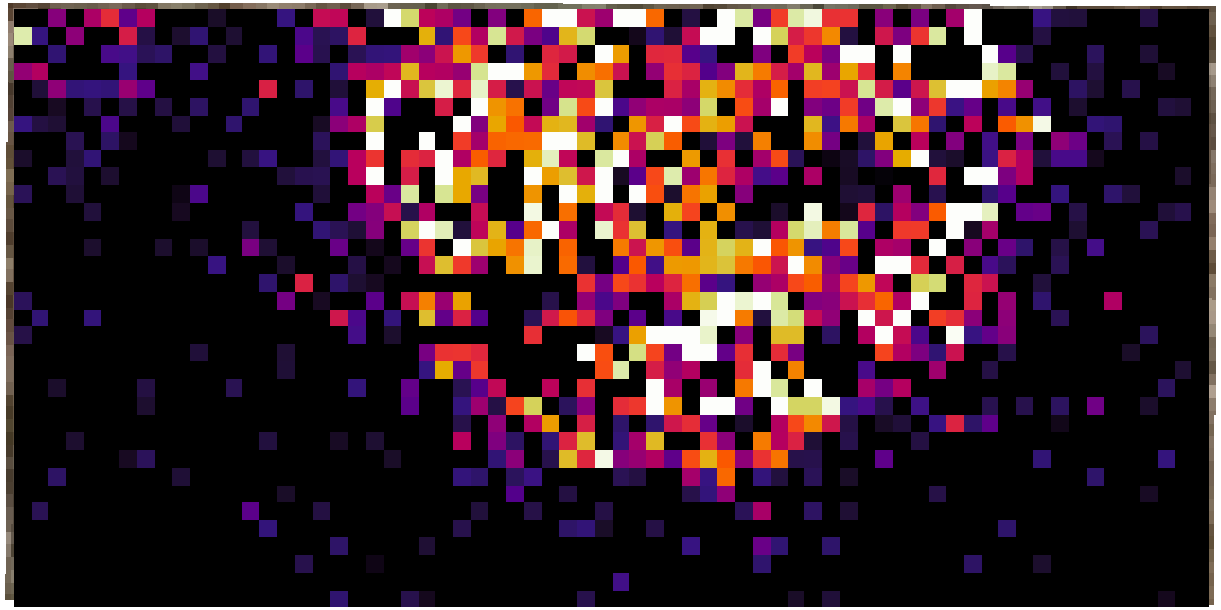

### Figure S1 (continued)

*m/z* 427.222

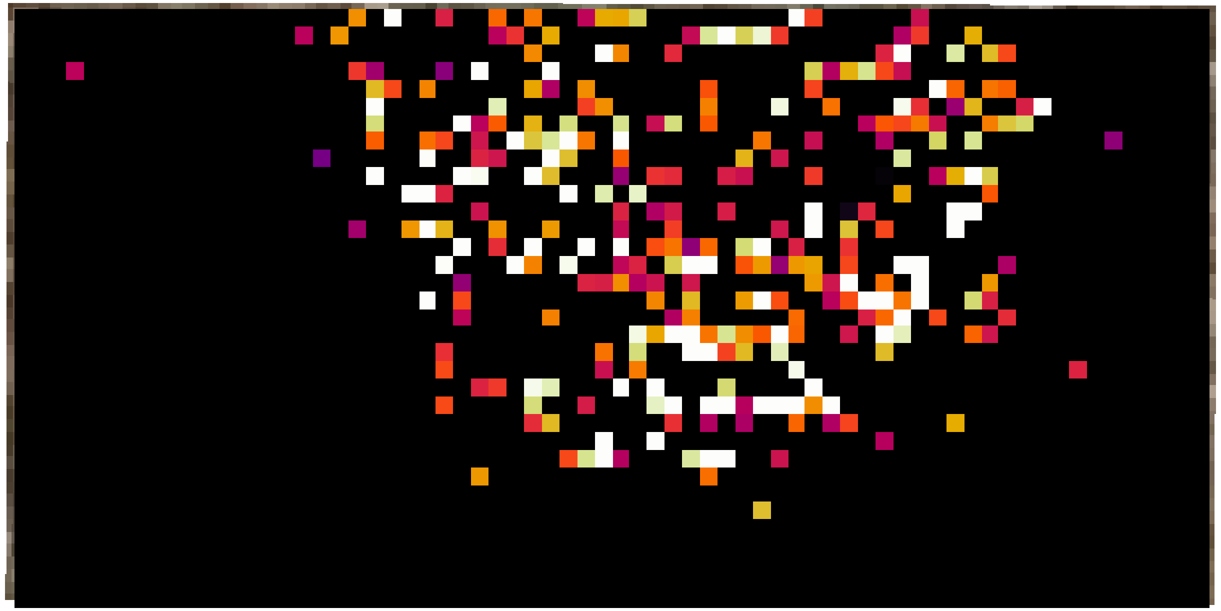

**Figure S1 (continued)**

*m/z* 471.212

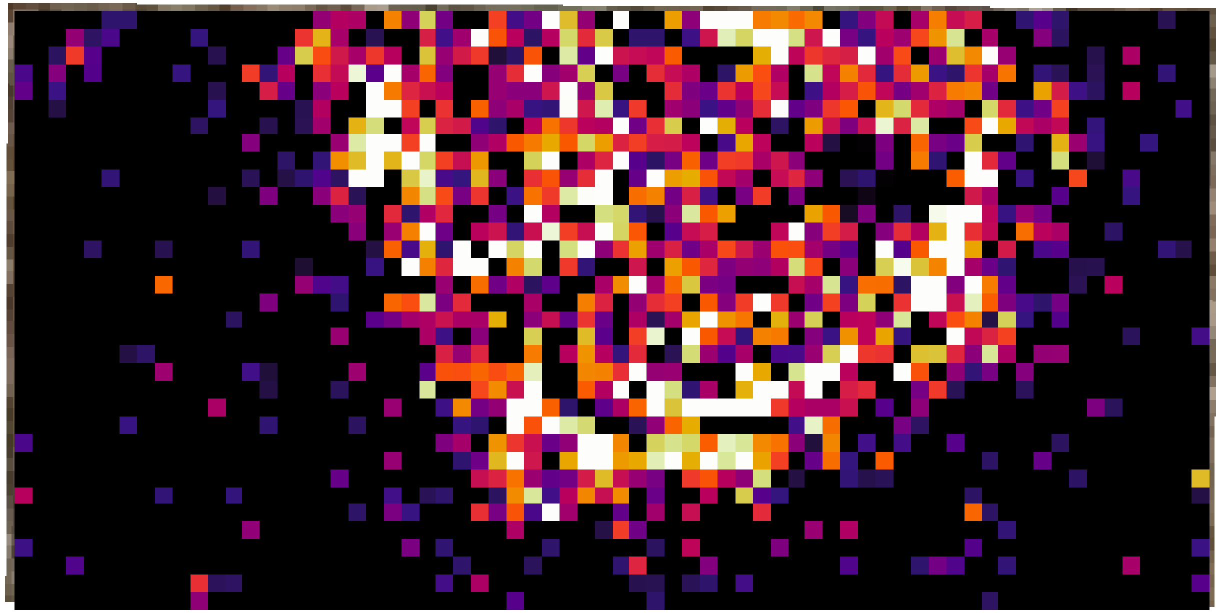
